# Single-antibody mutational scanning reveals poly-specificity and immune constraints in vaccination responses

**DOI:** 10.64898/2026.01.07.697678

**Authors:** Luca Johannes Schlotheuber, Simon Nørregaard Agersnap, Deborah Greis, Alessandro Streuli, Yohei Yamauchi, Klaus Eyer

## Abstract

Current repertoire technologies with single-antibody resolution usually assess binding to only one or a few recombinant antigens, neglecting variant diversity of antigens and poly-specificity of antibodies. Here, we introduce a method for analyzing antibody repertoires at single-antibody resolution against a large library of SARS-CoV-2 receptor-binding domain (RBD) variants, enabling the identification and analysis of recognized RBD variants by rare, poly-specific antibodies. This technique provided unique insights into single-antibody binding and escape profiles within a murine immunization model with different RBDs, offering valuable data useful to optimize vaccine design against emerging variants and study antibody poly-specificity.

## Introduction

SARS-CoV-2 is a RNA virus that continues to evolve into variants with increased virulence, immune evasion, and altered receptor binding^1–4^. The spike protein, especially its RBD, has been a key target for vaccine and antibody development^5–7^, but frequent mutations in the RBD reduced the effectiveness of highly specific antibodies and vaccinations over time, posing ongoing challenges to therapy and protection^5,6,8^.

To this end, techniques like deep mutational scanning (DMS) have transformed our ability to study sequence-function relationships by introducing and analyzing a wide range of amino acid (AA) changes across a protein^9,10^. Indeed, DMS enabled a detailed mapping of how mutations affect antibody binding, immune escape, virus infection, receptor binding or protein expression overall, and also informed vaccine design by identifying critical epitopes^11,12^. Yet, DMS is typically applied to a few antibody clones for characterization after their selection, or serum samples that consist of various antibodies, and may obscure rare and broadly cross-specific antibodies present in the full polyclonal repertoire.

To resolve these complex repertoires on the single-antibody level, droplet microfluidics and other advanced technologies have been developed to identify rare, high-affinity antibodies^15,16^ against only one or a few antigens^16–19^, and don’t generate data useful for DMS. However, these variant-specific binding properties are crucial in the context of viral evolution and could inform the development of broader-spectrum antibody therapies or vaccine^20,21^. Therefore, integrating single-antibody analysis with DMS could provide an unprecedented view into antibody-antigen specificity, enhancing our understanding of immune escape and cross-protective immunity.

## Results

To combine microfluidic single-antibody experiments with DMS-suitable data, we integrated an antigen library of SARS-CoV-2 RBDs (AA 331 to 531), taken from Starr *et al*.^10,24^, for the antibodies to be analyzed against. The mutational library contained an original reported diversity of 180,000 variants (Figure 1A). These RBD variants were packaged into T7 phages, amplified and the correct insertion as well as the mutation frequency of the DMS library verified by PCR and sequencing (Supporting Figure 1). After several purification steps, we obtained a phage suspension of around 10^11^-10^14^ pfu/ml for encapsulation and performed Sanger sequencing to confirm the correct orientation of the library and deep sequencing to confirm the number of variants per read and overall library diversity (Supporting Figure 2). Our obtained library translated into the presence of ∼50-80,000 phages per droplet. With roughly 10 protein copies/phage^25,26^, we estimated a total in-droplet RBD concentration of 30-50 nM, similar to other droplet assays used for antibody screening^15,16^, although individual variant concentrations are much lower due to their diversity.

**Figure 1.**
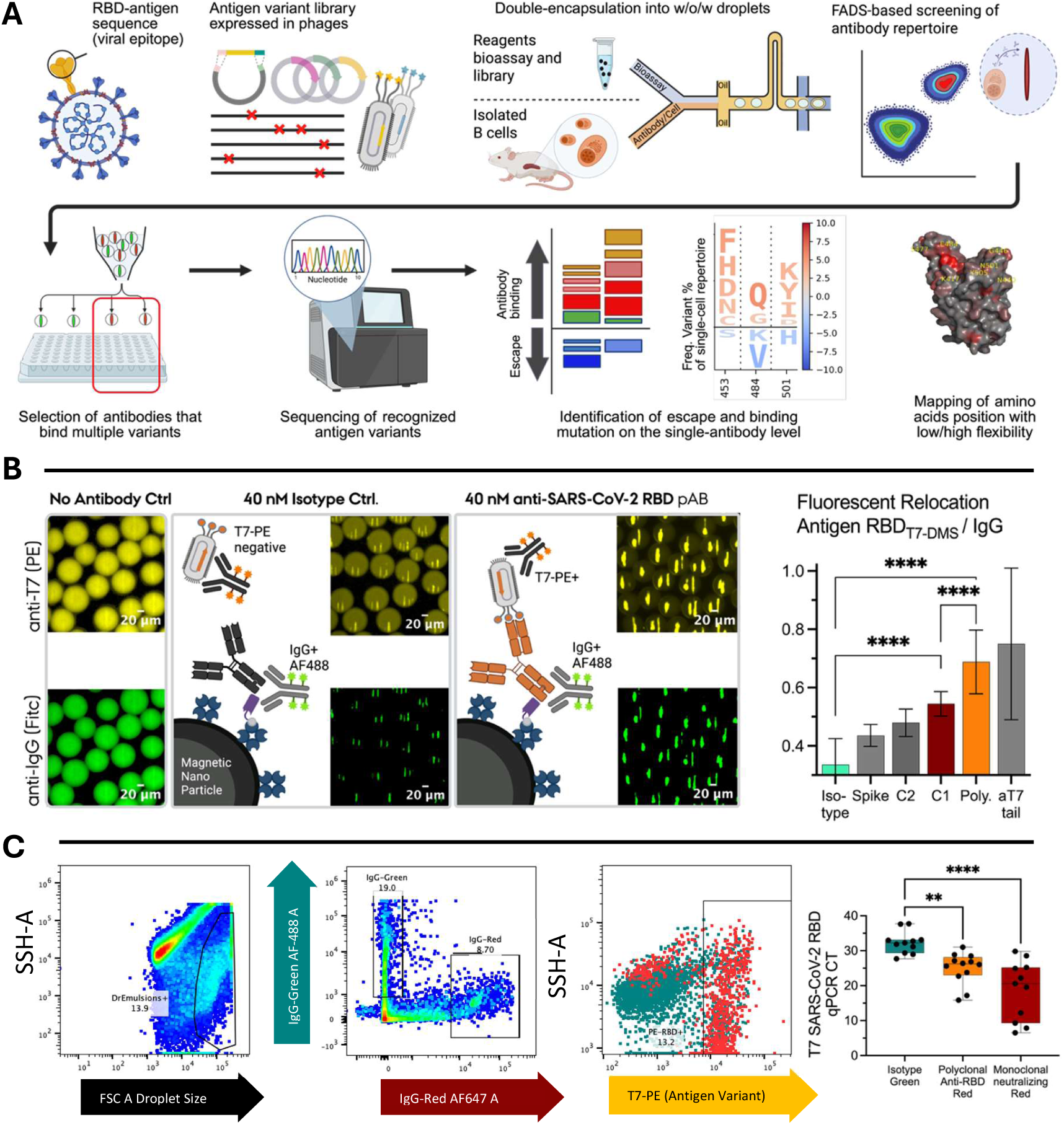
Single-poly-specific antibody analysis workflow. **A) Overview of the developed DMS workflow.** The RBD variant library was expressed in T7 phages and encapsulated in droplets along with a bioassay to capture and detect IgG. Droplets containing individual B cells were sorted for IgG secretion and RBD poly-specificity, the recognized antigens sequenced, and the data analyzed for antibody enrichment and escape. **B) Assay development and calibration of RBD-variant binding.** Left: the micrographs and scheme show the assay principle in the presence of no antibody, non-RBD-specific and RBD-specific antibody (pAb, both at 40 nM). Right: Relocation calibration experiments of the anti-T7-PE probe with different mAb and pAb antibodies at 40 nM. **C) FADS gating calibration** (left), and RBD-specific qPCR quantification of isolated variants post-sorting (right). Statistical analysis by one-way ANOVA, *p*< 0.05 (*), *p*< 0.01 (**), *p*< 0.001 (***), and *p*< 0.0001 (****).

We hypothesized that such assay conditions would favor antibodies recognizing a broader range of variants, with poly-specific clones yielding stronger binding signals. To test this hypothesis, we assayed the fluorescent signal using RBD-specific mono- (mAb) and polyclonal antibodies (pAb, SI Table 1), the latter as a proxy for poly-specificity. The broader specificity profile of the pAb was confirmed by ELISA (Supporting Figure 3). In droplets, RBD-specific mAbs were distinguishable from non-specific antibodies based on their binding signals (Figure 1B, Supporting Figure 4). Most importantly, pAb showed higher signals than the RBD-specific mAbs, despite higher affinities in mAb (Supporting Table 2), and was similar to the anti-T7 antibody control. Therefore, we concluded that the signal in our assay might “favor/bias” for recognizing a spectrum of RBD variants.

Next, as we aimed to analyze individual antibody clones secreted from cells, we implemented the strategy developed by Brower *et al*.^27^ to sort individual calibration droplets into discrete wells using fluorescence-activated droplet sorting (FADS). Here, we used the pAb, a neutralizing RBD-specific clone (nAb) and a non-RBD-specific mAb (negative control). To distinguish both populations (specific *vs*. non-specific) in mixed droplet populations, we included two different fluorescently labeled IgG detection probes to identify both populations and their overlap in antigen binding. In flow, the RBD signal could be differentiated, with an in-gate specificity of ∼87% and at a sensitivity of ∼82% (Supporting Figure 6). The conditions were also distinguishable in qPCR, amplified from individual sorted droplets (Figure 1C), further correlating with concentration of RBD-specific antibody (Supporting Figure 5).

Looking further into the PCR-amplified sequences after sorting, we determined the efficiency of successful PCR amplification to be around 38% from droplets containing RBD-specific antibodies. However, amplification was very specific, as <2% of droplets containing a non-specific antibody led to RBD sequences being amplified (Supporting Figure 7). Even though our calibration suggested that not all possible amino acid variants of the RBD may be present in each droplet, our repertoire data indicated that many known RBD variants existed and pAb droplet showed overlapping, reproducing signatures across the sampled repertoire (Figure 2, Supporting Figure 11). Next, we immunized BALB/c mice either with SARS-CoV-2 RBD (wt) or B.1.351 RBD (B.1.351, mutations K417N, E484K and N501Y, Figure 2A) to induce distinct antibody repertoires. We first characterized the induced antibodies using stationary droplet microfluidics on different days, and decided on 7-day post-secondary immunization where the combination of frequency of secreting cells, their secretion rates and affinity profile were deemed optimal (Supporting Figure 8)^15^. Most importantly, when measured against the RBD variant library inside droplet arrays, a fraction of cells displayed high T7 binding, signals exceeding the calibration signal for pAb (Figure 2A, Supporting Figure 4). This indicated the potential presence of poly-specific RBD-binding IgGs, especially in B.1.1.35 immunized mice (Figure 2A). Serum-based analysis against the variant phage library further hinted towards the recognition of a broader spectrum of RBD variants in the B.1.1.35 condition (Supporting Figure 3C).

**Figure 2.**
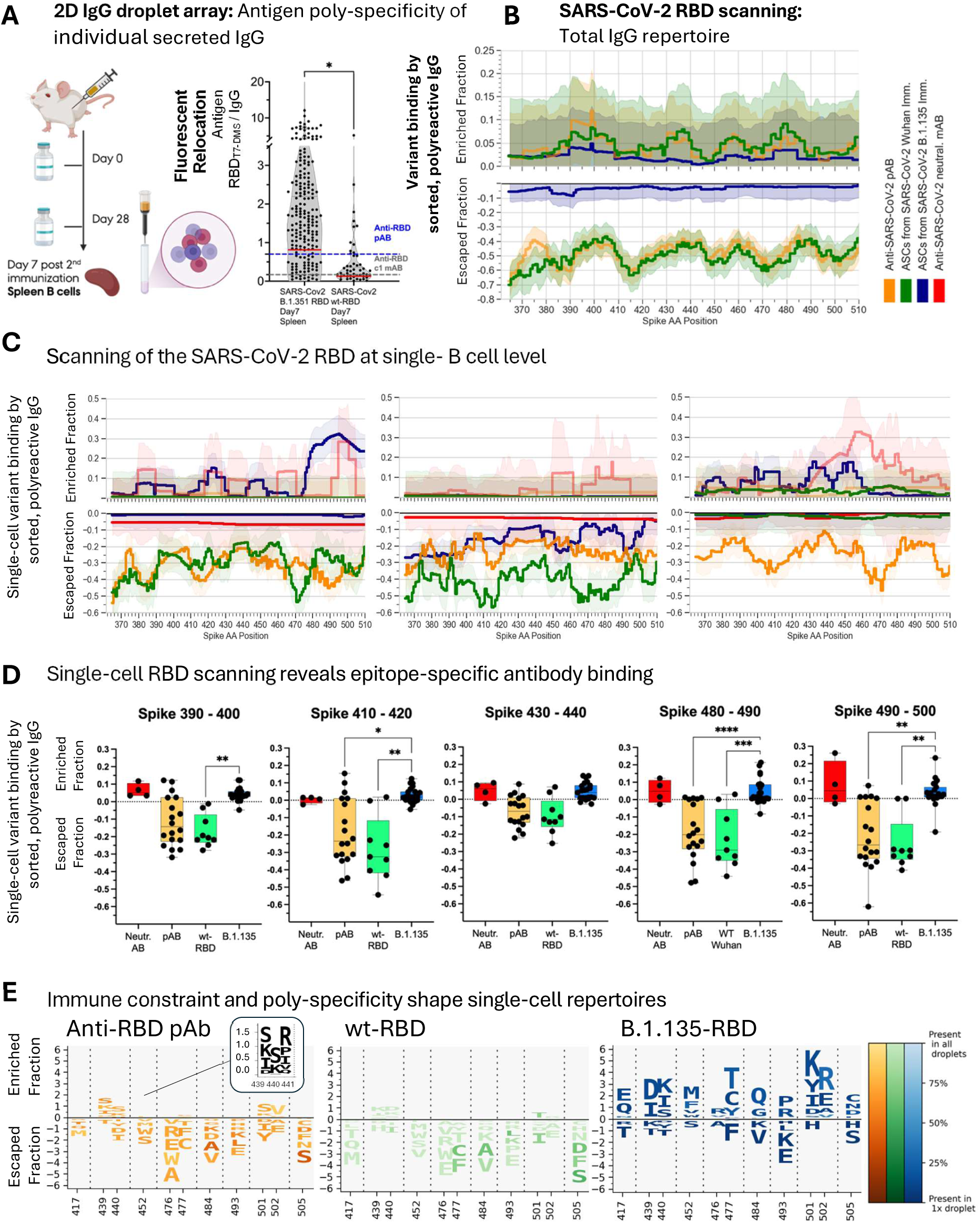
DMS results of summarized and individual RBD-poly-specific IgGs of immunization-induced repertoires. **A)** Immunization (left) and RBD variant binding ratios extracted from stationary droplet arrays (right), showing antigen relocation (anti-T7 tag) relative to the amount of secreted IgG from individual B cells (blue line – median, dotted– threshold pAb from calibration). **B-E)** Variant binding (change in frequency of mutation) of antibodies from pAb (orange), wt-RBD (green) and B.1.351 immunization (blue) **B**) from all IgG+ poly-specific antibodies (Enriched and escaped fraction) from samples with at least 5 sequenced droplets; shades represent 95% confidence interval **C)** of three representative individual droplet measurements from each antibody repertoire (separated into binding and escape fraction), and**D**) of each single-cell repertoire compared across specific RBD regions. **E**) (Selected) mutations in the RBD within the repertoires (DMS Logo plot, height proportional to binding ratio E_R_). Computed binding shifts in **B-E** show the median (Log10 transformed) binding ratio E_R_ of mutations at a given position. For statistical comparison (**D**) E_R_ binding ratios were binned across regions; boxes mark the 25th/75th percentile with each dot representing the median Log10 binding ratio of a sampled droplet. Statistics performed by one-way ANOVA of ranks by Kruskal-Wallis (KS) test, *p*< 0.05 (*), *p*<0.01 (**), *p*< 0.001 (***), and *p*< 0.0001 (****). Total number of (single antibody) barcodes in the analysis: n=66.

Next, we examined the recognized antigen variants to compare general and individual antibody poly-specificity between the two immunizations. We sorted individual droplets with high T7-RBD relocation and sequenced the recognized variants. Indeed, the median Ab binding frequencies revealed immunization-specific binding profiles with distinct clusters in the RBD at 390-410 and >470, inside known antibody binding and neutralization sites (Figure 2B)^28^. Notably, when split by whether mutations increased or decreased IgG binding to RBD variants, the B.1.135-RBD immunization induced an overall poly-specific repertoire that recognized these sites more frequently and with higher binding rates than the wt-RBD, which, as expected, closely matched the pAb profile (Figure 2B/C).

At the single-antibody level, more distinct profiles of RBD variant binding were visible (Figure 2C). Secreted IgGs from wt-RBD–immunized mice rarely targeted AA 480–510, but in contrast, multiple clones extracted from B.1.135-RBD-immunized mice displayed antibodies with shifts towards increased binding in those regions. Overall, B.1.135 derived antibodies showed surprising heterogeneity, with some clones showing preferences to 470-500, possibly interfering with ACE2 binding (Figure 2C, left), while others showed shifts, indicating potential immune escape (Figure 2C, center). Finally, we could identify few clones binding to less commonly targeted sites (AA 430-450, Figure 2C, right).

To further identify and compare immunization specific signatures across the single-antibody repertoire, we compared binding across five specific binding regions of the RBD (Figure 2D). Although we observed large clonal binding diversity before in single-cell plots, trends towards certain epitopes were observed across the immunization conditions: For instance, no significant shifts were detected at position 430 between repertoires, while regions 410–420 and 480–500 showed substantially different binding patterns in B.1.135-RBD induced antibodies compared to pAb and wt-RBD conditions. PCA further clustered the data according to immunization, showing strong overlap between wt-RBD and pAb repertoires, while B.1.135-induced B cells formed distinct clusters (Supporting Figure 12-13).

To further explore mutation-specific effects at the single-cell level, we generated logo plots showing binding and escape ratios for certain, relevant RBD AA positions, with color intensity reflecting the recognized prevalence of mutation across the single-droplet repertoire (Figure 2D, 2E). Here, B.1.135-RBD immunization produced IgGs with high RBD binding ratios, >5-fold higher than those elicited by wt-RBD (Figure 2C/E) and yielded a highly diverse antibody repertoire. For some amino acid mutations, these binding shifts at the individual antibody level are occurring with more than 50-fold increase upon binding (Log10 of 1.7) relative to the DMS library (Figure 2E), further underlying the single cell heterogeneity. Interestingly, while antibodies from B.1.135-immunized mice displayed several IgG with high and distinct RBD binding, in contrast pAb showed low binding ratios, but across many mutations (>7 AA, Figure 2E, enlarged).

This suggested that the pAb and wt-RBD derived poly-specific antibodies show preference to the Wuhan variant (wt) as expected, but also may more frequently bind conserved, overlapping epitopes (Figure 2B/E). Most notably, the B.1.135-RBD induced poly-specific antibodies could recognize key escape mutations such as N501Y, absent in wt repertoires, and displayed local enrichment in known class I-III binding regions^29^, including the receptor-binding motif (RBM), suggesting potential competitive neutralization behavior (Figure 2E, Supporting Figure 14). Conversely, some major escape mutations, specifically E484K, were rarely targeted in any of our single-cell repertoires, despite their poly-reactivity, even by B.1.135-derived antibodies, indicating a potential shared mechanism of immune constraint. Within the escaped fraction, few, “large” gaps in binding exist, pointing to mutations which decrease IgG binding across the repertoire, many shared with wt-RBD (Figure 2E, Supporting Figure 9).

## Discussion

In this work, we introduced an antigen scanning approach with single-antibody resolution to investigate SARS-CoV-2 antibody responses, focusing on the sum of and individual poly-specific antibodies. As a proof-of-principle, we compared antibody repertoires after immunization using recombinant wt-/B.1.135-RBD. In our experiments, B.1.135-RBD induced poly-specific IgG-secreting cells with broader antigen diversity, detected by fluorescent relocalization and sequencing of recognized RBD-expressing T7 bacteriophage variants. Additionally, the sequenced antigen libraries from individual mAb from B.1.135-RBD-immunized mice showed an increased shift in RBD binding, evidenced by unique mutation binding profiles and their clonal variability.

Our data showed that only 1:10-1’000 RBD-specific antibodies exhibited such high variant binding, which may be overlooked in bulk serum analysis. Notably, some mutations, such as 484K, were difficult to target by poly-specific antibodies from either immunization. This limitation was not due to poor overall immunogenicity, as B.1.135 RBD immunization generated high ELISA titers and successfully targeted other known escape mutations like 501Y. Interestingly, several isolated RBD-specific clones exhibited high poly-variant specificity, including recognition of key variants 417N and 501Y, and mutational scanning further revealed a spectrum of antibodies targeting different regions of the RBD, allowing classification into one or more SARS-CoV-2 antibody classes I-IV based on structural mapping.

Interestingly, many of our findings based on poly-specific antibodies were consistent with previously reported immunogenicity differences between lineages^30,31^, although here, they were resolved into individual poly- and non-poly specific antibodies. Overall, single-cell antigen scanning offered advantages over serum analysis, but achieving sufficient and uniform sequencing and sampling depth across samples remained a challenge. Increased sequencing depth may allow for precise RBD epitope mapping with high-resolution in the future. Furthermore, the success rate of useful data extraction from the IgG secretion event to retrieving the antigen sequence was limited, but when data were extractable, they were very specific.

In conclusion, our scanning approach offered a quantitative and qualitative method to dissect the highly diverse binding profiles of antibodies at single-cell level, allowing to compare different immunizations for their poly-specific antibody repertoire. Coupled to antibody sequencing, these poly-specific antibodies further hold promise as therapeutic candidates, especially when integrated with viral evolution predictions, and might characterize immunizations in much more depth.

## Acknowledgements

The authors gratefully acknowledge Cyril Brunner from Bruker Scientific Technology Co., Ltd. for making their instruments available and conducting experiments for SPR analysis. We also thank the Flow Cytometry Core Facility at ETH Zurich, particularly Dr. Florian Mayr, for droplet sorting and support with flow cytometric analysis. We further thank the Functional Genomics Center Zurich, and particularly Dr. Qin Zhang, for the support on library preparation and sequencing. The European Research Council starting grant (Grant #803′363) and the Swiss National Science Foundation (Grant #310030_197619) supported this work.

## Conflict of interest

All authors declare no conflict of interest.

## Methods

### Generation of the SARS-CoV-2 protein-expressing library

Antigen libraries were generated according to Starr *et al*.^10^ by iterative rounds of low-cycle PCR with pools of mutagenic synthetic oligonucleotides that each contain a randomized NNN triplet at specific codon sites. SARS-CoV-2 Spike Receptor Binding Domain Deep Mutational Scanning Library was a gift from Jesse Bloom (Addgene #1000000172). Libraries were cloned into a T7 vector arms using primers outside the RBD-DMS and barcode sequence (Supporting Figure 1). The PCR product (1,490 bp) was size selected from 2% TAE, purified, and dual digested using EcoRI and HindIII. The RBD variant expressing phage library was then cloned and expressed using the T7 Select kit (Novagen, Merck Millipore). In short, the purified DNA fragments were cloned into T7 vector arms using T4 DNA Ligase (ThermoFisher) at a 3:1 molar ratio of insert DNA to vector arm DNA according to manufacturer’s instructions^26,32^. The use of restriction digest rather than PCR eliminates the possibility of PCR strand exchange, possibly resulting in misfolded RBD variants. This resulted in a 1,045 bp fragment, the length confirmed through gel electrophoresis (ThermoFisher Taq PCR Master Mix 2x) which was then ligated into the predigested T7 vector arms (Supporting Figure 1).

### Library phage packaging, propagation and storage

First, 5 µl of the ligated T7 vector (1 µg) was incubated with 25 µl T7 packaging extract according to the manufacturer’s instruction (Novagen, Merck Millipore) for 2 h at RT for *in vitro* packaging of the phages. The reaction was stopped by adding 270 µl of LB medium. For T7 pre-stock propagation, a 50 ml LB medium containing 100 µg/ml ampicillin was inoculated with 10 µl of bacterial glycerol stock, either BLT5403 or BLT5615 (Novagen). Once the pre-stock had grown to OD_600_ of 0.8 at 37°C, the culture was infected with 50 µl of the *in vitro* packaging reaction. The culture was incubated further while shaking at 37°C until signs of bacterial cell lysis were observed (reduction in OD and visible accumulation of white debris). Initial T7 pre-stock was centrifuged at 4,500 g for 10 min and used to infect larger stocks (400 ml) directly or frozen down in Glycerol for later use.

### Initial quality control of phage library

Successful ligation and propagation were confirmed by PCR on T7 DNA after amplification in *E. coli* with RBD insert specific primers and primers amplifying the multiple cloning site (MCS) of the T7 vector including any DNA inserts (RBD short primer, T7 UP/DOWN primer, Supporting Table 1, Supporting Figure 1A). The RBD phage sample showed bands at the expected lengths of around 648 bp for the RBD insert-specific primer and 1,149 bp (1,045 bp insert + 144 bp MCS) for the T7 MCS primers (Supporting Figure 1). To further control the presence of phages not containing RBD sequences, a control plasmid without an insert for protein display was provided in the kit and packaged and grown in *E. coli*. The control and RBD phage samples both displayed a small amplicon at 144 bp when using the primers targeting the Multiple cloning site (MSC), indicating that the RBD Phage library sample also contained some phages not containing a RBD insert. Correct cloning was further confirmed by Sanger sequencing (Supporting Figure 1B).

### Library phage purification

The obtained lysate was purified and the phages concentrated in three steps. First, the lysate was centrifuged at 8,000 g for 10 min to remove cellular debris, and the supernatant was transferred into a sterile bottle, and adjusted to 0.1 M NaCl by the addition of 5 M NaCl. Second, the supernatant lysate was mixed 1-to-6 with a 50% (w/v) PEG8,000 solution (Carl Roth) and incubated at 4°C for 24 h to precipitate the phages. After centrifugation for 10 min at 7,000 g and 4°C, the phages were collected as a pellet. The supernatant was discarded, and the T7 phages were resuspended in 1.2 ml consisting of 1 M NaCl, 10 mM Tris-HCl, 1 mM EDTA at pH 8 (TE buffer). Thirdly, CsCl-TE buffer density gradient ultracentrifugation was employed for further purification following the T7SelectR manual (Novagen, Merck Millipore). The PEG-purified T7 phages were added on top of the CsCl density gradient, and ultracentrifugation was performed in the Optima XPN (Beckmann Coulter, SW41 rotor, 35,000 rpm, 60 min, RT). Blueish T7 phage bands were collected from the respective interface (2:1 CsCL:TE) by using a 22–25 Gauge, 25 mm long needle attached to a syringe. To desalt the generated T7 antigen libraries, the T7 phages were purified using Slide-A-Lyzer™ MINI Dialysis Device, 7K MWCO or snakeskin dialysis tubing, 3,5 MWCO (both Thermo Fisher) overnight at 4°C against B cell buffer media (cell buffer). Bacteriophage T7 displaying the antigen library were stored short term at 4°C in cell buffer, and in 20% glycerol buffer at -80°C for longer times.

### Plaque assay, qPCR RBD specific C_t_ and its correlation

First, to determine the phage titer after purification, a plaque assay was performed as described in the T7SelectR system manual (Novagen, Merck Millipore). Briefly, *E. coli* host cells were grown as described above to an OD_600_ of 1. A 1:10 dilution series of the phage stock was generated in LB + Ampicillin (100 µg/ml) medium. To 250 µl of the host cell suspension, 100 µl of the phage dilutions were added to a 15 ml reaction tube, and 3 ml top agarose was added, mixed through pipetting, and spread evenly on a prewarmed (37°C) LB agar plate. These plates were prepared in advance by pouring 25 ml liquid LB agar into 100 mm Petri dishes and stored at 4°C after setting the agar. The plates were inverted once the agarose had set and incubated at 37°C for 4 h. The phage titer was calculated using the number of counted phage plaques (plaque forming units, pfu) in the bacterial lawn on a given plate and the corresponding dilution factor. The pfu was used to correlate the C_t_ value measured by qPCR in duplicates. Two 1:10 dilution series of purified RBD library phage stock were prepared and analyzed through qPCR to relate the C_t_ values to the titer measured by plaque assay according to the equation below (Supporting Figure 1C). Due to the simplicity of qPCR, this method was used in subsequent preparations and dilutions to estimate the phage titer in generated phage solutions:

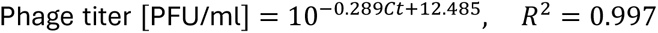

### Preparation of nanoparticles for antibody capture and detection

Nanoparticles were functionalized as previously described^15,33^. In short, 300 nm-sized magnetic streptavidin nanoparticles (Ademtech) were functionalized with CaptureSelect *V_H_H* Biotin Anti-LC-kappa (ThermoFisher), and 140 nM of anti-IgG-Fc Alexa 647 or 488 detection probe (SouthernBiotech, see also Supporting Table 1 for more information) was added to a final droplet concentration of 70 nM. The phage library was added to the particles, presenting RBD, at a final in-droplet concentration of around 10^5^ phages/droplet. For the detection of T7 phages and their relocation to the beads, we used a goat anti-T7-Tag PE antibody (Abcam, 4 nM final in-droplet concentration).

### Immunization and preparation of murine B cells for analysis

Formulations for immunizations were prepared by adding 10 μg/100 μl of the respective recombinant RBD in PBS/Alum (AcroBio, mixed 1:1, Supporting Table 1) for 2 h at RT. These experiments were performed according to ARRIVE and institutional guidelines, local and Swiss Federal regulations and were approved by the cantonal veterinary office Zurich (license ZH215/19). BALB/c mice (Janvier Labs, female, age 8-10 weeks at start) were immunized intraperitoneal with 10 μg RBD twice four weeks apart. At indicated days after secondary immunization, spleen and bone marrow were collected. To isolate murine B cells from the spleens, the spleens were disintegrated using a 40 μm cell strainer and plunger. The cells were washed three times in buffer (PBS, +10 mM EDTA) and filtered again (40 μm). The cell suspension was treated with BD Pharm Lyse (BD Biosciences) for 5 min at RT. Afterward, the cell suspensions were diluted immediately (PBS, +10 mM EDTA) and filtered again through a 40 μm cell strainer. The single-cell suspensions were centrifuged at 4°C, 400 g for 5 min, and washed in buffer twice before using the murine Pan B cell isolation kit and MultiMACS Cell24 Separator Plus (Miltenyi Biotec) according to the manufacturer’s instructions. The untouched cells from the B cell lineage were collected and counted. The cells were afterward stained with CellTrace Violet (Thermo Fisher) for 10-15 min on ice, subsequently washed with RPMI with 10% FBS, and collected by centrifugation. Finally, the cells were resuspended in RPMI 1640 without phenol red (Sigma-Aldrich), supplemented with 5% KnockOut Serum Replacement (Thermo Fisher), 0.5% recombinant human serum albumin (Sigma-Aldrich), 25 mM HEPES (Thermo Fisher) and 0.1% Pluronic F-127 (Thermo Fisher) to achieve cell-to-droplet ratios between 0.2 and 0.4 after encapsulation according to previous work^33^.

### Preparation of calibration antibodies for in-droplet measurements

The calibration antibodies used are listed in Supporting Table 1. The antibodies were diluted in cell buffer and were used for droplet generation instead of the cells. For calibrations, specific antibodies and particles were encapsulated in distinct inlets to prevent cross-linking prior to encapsulation.

### Droplet chip and chamber fabrication, generation of droplets

The chips, chambers, and emulsions for stationary droplet arrays were produced as previously described^15,34,35^. For sorting experiments, we employed the protocol as outlined by Brower *et al.*^27^ with slight modifications. Here, the original design for the microfluidic double emulsion chip was slightly altered and optimized for the use of the nanoparticles by implementing a funnel-shaped encapsulation flow and resizing the encapsulation nozzle (see Supporting Figure 15). We used as a 1^st^ continuous phase 2% 008-Fluorosufactant (RAN Biotechnologies) in HFE-7500 fluorinated oil (3M), and PBS with 1% Tween-20 (SigmaAldrich) and 2% Pluronic™ F-127 (ThermoFisher) as 2^nd^ continuous phase. Droplets were generated with flow rate ratios of 150/150/600/6000 µl/h, respectively, and collected in low-binding 1.5 ml tubes (Eppendorf).

### Image acquisition and analysis of stationary droplet arrays

For calibration, calculation of antibody secretion and antigen binding, fluorescence imaging of stationary droplets was performed using a Nikon TI2 Eclipse epifluorescence microscope. The droplets were maintained in a darkened cage incubator (Okolab) at 37°C within a custom-made observation chamber^15,34,35^. Imaging was conducted with a 10× objective (NA 0.45), and channel-specific camera settings were applied (Orca Fusion or R2, Hamamatsu). Emission was detected with appropriate band-pass filter sets (DAPI, FITC, TRITC, Cy5; all Semrock) and illuminated using a SOLA LED light source (Lumencor). At each time point, a large microarray of images was captured, with all channels recorded sequentially for each image before the stage advanced to the next position. Measurements were taken every 15 or 30 minutes for 10 time points for experiments containing cells, and once for calibration experiments.

### Droplet breaking and RT-PCR

For correlating and calibration antibody concentrations and qPCR of T7-antigen after sorting, 100 µl of single droplet emulsions were generated, collected in a separate tube, 1 ml of PBS was added, and the emulsion was demulsified by the addition of 15 µl perfluoro-1-octanol (Sigma-Aldrich). Next, the solution was mixed and filtered through a 70 µm cell strainer. The nanoparticles with attached T7 phages were centrifuged at 500 g for 5 min and the supernatant was discarded, and the particles were further washed twice with PBS with added 0.5% Tween-20 (Carl Roth) and twice with PBS with added 10 mM EDTA. To extract phage DNA from the T7 phages attached to the beads, the nanoparticles were resuspended in 20 µl PBS with added 10 mM EDTA, and incubated at 65°C for 10 min, centrifuged at 4,000 g, 4°C for 5 min, and kept on ice. For qPCR, 2 µl of the samples were mixed with PowerTrack SYBR Green Mastermix (Thermo Fisher) in 10-20 µl per reaction. All reagents were mixed on ice with a final primer concentration of 500 nM, and the reaction was carried out in 384-well format in Quantstudio™ 7 Flex Real-time qPCR system (ThermoFisher) in standard cycling mode for 40 cycles.

### T7 library bio-panning ELISA

To characterize the spectrum of variants bound by the used calibration antibodies, we established a bio-panning ELISA method. To do so, T7 variant libraries were immobilized on Maxisorp Nunc-Immuno 96-well plates (Thermo Scientific), and the plates were incubated for the coating of the library overnight. Next, the plates were washed three times with PBS containing 2 mM EDTA and subsequently blocked with RPMI + 10% FBS + 10 mM EDTA for 2 h. After blocking, the plates were washed and incubated with serial dilutions of the respective antibodies for 1 h at RT. Plates were washed again twice with PBS containing BSA, followed by the addition of rabbit anti-mouse HRP (50 ng/mL, A27025, ThermoFisher Scientific), each diluted to 50 ng/ml in PBS with BSA (100 μl per well). Afterward, the plates were washed three times with PBS. For fluorescence measurement, fresh detection reagent was prepared by mixing PBS with HSA, 30 % H₂O₂ (final concentration 100 μM), and Amplex Red (final concentration 5 μM, Invitrogen A22177), and 100 μl was added per well. Directly afterward, the fluorescence signal (excitation at 535 nm, emission at 590 nm) was recorded using a plate reader (Tecan infinite m200).

### Total IgG ELISA of serum

Mouse sera were diluted ranging from 1:50 to 1:50,000. 200 µL of each sample was loaded in duplicates in wells of microtiter plates (Multiscreen high binding F-bottom 96 well clear polystyrene, Millipore Corporation) and incubated over night at 4° C. Plates were blocked with 5% BSA in PBS over night at 4° C. 100 µL rabbit anti-mouse HRP (50 ng/mL, A27025, ThermoFisher) were added to the plates for 2 hours at room temperature. Binding was visualized with 100 µL TMB substrate for 30 minutes. Plates were washed after each step four times with washing buffer (PBS, pH 7.4 containing 10 mM EDTA). A mouse anti-CD20 IgG2a antibody (clone L26, antibodies.com) was used as positive control. Negative controls consisted of PBS. Reaction was stopped with 100 µL 0.16 M H_2_SO_4_ and absorbance was measured at 450 nm using plate reader (Tecan infinite m200). Coefficient of variation (CV) was determined for each duplicate, and samples with a value >15% were excluded.

### SPR measurements of calibration antibodies

SPR measurements were performed on a SPR-24 Pro from Bruker. Interaction analysis was performed using a Titration Cycle Kinetics (TCK) mode. Full association phases were recorded for all analyte concentrations, while a full dissociation phase was measured only for the highest concentration. Antibodies were captured onto a protein A/G sensor surface at a flow rate of 5 µL min⁻¹ for 240 s, achieving capture levels of 618 RU (Ab c1), 1100 RU (Ab c2), 533 RU (Ab c3), 841 RU (Ab N), 1090 RU (Ab P), and 987 RU (RaM). The full-length SARS-CoV-2 Spike protein was subsequently injected at a flow rate of 30 µL min⁻¹ with a contact time of 120 s and a dissociation time of 1800 s. Measurements were performed as sextuplicates. Data was analyzed in Analyzer 4 (Bruker Daltonics) and final plots were generated in Prism (GraphPad). Sensograms were recorded for six analyte concentrations.

### Double emulsion-based measurement and sorting

For the sorting of the droplets, the double-emulsion droplets were collected and incubated for 1 h at 37°C to stimulate antibody secretion. Afterward, the double-emulsion droplets were sorted using either an Aria II Cell Sorter (BD) or a ThermoFisher BigFoot with a nozzle size of 80 µm. The gating strategy for calibration antibodies is shown in Figure 1C and in Supporting Figure 6. The events of interest containing the droplet and the cell were sorted into individual wells of a 96-well plate (ThermoFisher PCR plate, fully skirted). Droplets were always encapsulated with a Goat IgG detection probe (Southern Biotech), either against Mouse IgG or Human IgG. For calibration experiments the same detection probe with AF647 and AF488 was used to discriminate droplet populations between positive and isotype. For gating, an initial threshold was applied on forward scatter (FSC-A) to exclude small particles and oil debris from the subsequent analysis. Next, a gate was set to identify larger double emulsions. Subsequent positive events where either identified using the IgG+ and anti-T7-Probe for antibodies or for B cells using, in addition, a cell stain (DAPI). After sorting, the 96-well plates with sorted droplets or control samples were washed twice on a magnetic stand to remove (left-over) unbound phages, resuspended in PBS + 10 mM EDTA, incubated at 60° C for 5 min, centrifuged, and stored at -20°C until PCR.

### PCR and amplicon sequencing of T7-antigen sequences

The PCR protocol followed instructions for Illumina Nextera (Nextera DNA Library Prep Reference Guide, Illumina). PCR was performed using high-fidelity Phusion Polymerase (New England Biolabs) in two steps. First, for each well (originally containing a single droplet) primers binding to T7 vector arms (containing Amplicon overhang UMI sequences) were used to amplify the RBD DMS insert. DNA quantity of amplicons was checked using Qubit DNA quantification kit. Each binding sample was amplified in a 2^nd^ PCR using a unique primer combination of N7XX and N5XX (barcoded) secondary primers containing i5/i7 illumina adapters (Microsynth). Amplicons were validated using the Tape station (Agilent) and a qPCR to check adapter content. For sequencing, the samples were pooled, the DNA concentration normalized and sequenced in a single run with the unaltered deep-mutational scanning library, separately barcoded. Library and barcodes were always sequenced in the same Illumina run to avoid comparing frequencies across technical, sequencing replicates. Data was run one paired-end NextSeq 2000 with 10% Phyx (Illumina), and the data was demultiplexed from FASTQ files using bcl-convert, filtered using FAST-QC, trimmed, merged using PAIR (paired-end read merger on Galaxy) for forward and reverse reads and finally aligned against the SARS-CoV-2 receptor-binding-domain Wuhan reference sequence using BWA-MEM2 in BAM format. From alignment files per-barcode amino-acid mutations and variants were quantified. A Jupyter Notebook “BarcodeCount” with a step-by-step count computation is available https://github.com/lucaSCHLTBER/SingleAntibodyDMS/.

### Analysis of single-antibody mutant library sequencing and computation of per-barcode/per-cell “binding” enrichment and “escape” fractions

For each (monoclonal) antibody (a given barcode from a sequenced droplet), we computed the enrichment, largely following the work of Fowler et al. ^36^ and Greany et al. ^37^. In short, we computed enrichment scores/ratios (E_R_) from Amino acid counts and their frequencies across all full-length reads. Counts and frequencies were stored separately for each amino acid of each single droplet barcode. For a given demultiplexed barcode we calculated the frequency F and the Enrichment Ratio E_R_ for a variant V relative to the variant frequency within the DMS library prior to antibody binding (F_lib_):

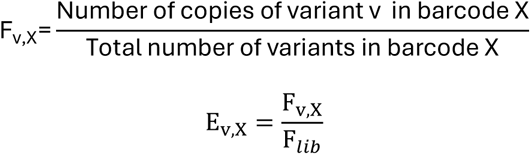

“Escape fractions” were defined as E_R_ values between 0 (meaning the variant always escapes antibody binding) and 1 (the variant is as frequent as in the original library population). “Binding fractions” are indicated by E_R_ > 1, meaning amino acids are preferentially bound by a specific antibody (increasing their frequency). An enrichment score (E_R_) of 10 indicates an antibody is increasing the variant frequency by 10x. We also applied a computational filter to remove variants with low sequencing counts or stop codon mutations that might cause escape of antibody binding simply by leading to poor expression of properly folded RBD on the T7 surface. We further excluded nucleotides from the very end and start of the RBD as we observed slightly reduced phred scores at the start and end of reads from MultiQC data. All sequences were analyzed and stored according to their barcode and mutation position relative to the Wuhan SPIKE protein sequence. We annotated each barcode according to its immunization or antibody source origin (murine secreting B cells from wt-RBD or B.1.135 RBD vaccinated mice, polyclonal antibody or anti-SARS-CoV-2 SPIKE neutralizing antibody).

### Identification of sites of poly-specificity, binding and escape

For line plots of all antibodies, the median E_R_ was calculated for all amino acids for each barcode separately. For the analysis (Figure 1, Figure 2) only non-synonymous mutations at each position were considered. For better comparison of enrichment (variants increased by antibody binding by 1x, 10x, 100x, 1000x), Log10 was computed for binding and escape fractions. For single-antibody line plots, a single barcode (cell or antibody) was randomly sampled from each immunization condition. To better assess poly-reactivity/specificity for a given spike RBD position, median of log10 ratios were computed across all amino acids. For Figure 2B, median was computed across all E_R_ from all IgG antibody profiles (without single-cell representation) of a given immune repertoire. For Figure 2B/2C, gaussian smoothing was performed on the enrichment ratios using a rolling window (window size = 10 AA residues) which averaged the median Log10 binding to reduce noise and enhance signal clarity around local epitopes. Line plots were underlined with a shaded region surrounding the average, showing the range (min-max) of smoothed E_R_ enrichment ratio values. In contrast, for amino acid logos (Figure 2E), within binding and escape fractions (log10 median) E_R_ was computed separately for each amino acid, but across all droplets of a given immunization condition. Color coding was chosen to visualize poly-specificity (how frequent an AA mutation was present among all antibodies of that repertoire). Dark colors indicate only few antibodies showed this specific median binding against that mutation, while fainter colors indicate that a higher proportion of antibodies contributed to a particular amino acid binding (median). DMSlogo^38^ package was used plot logo mutation profiles.

### Assessment of antibody profiles and variant diversity and similarity

To compare variants sequenced from single droplet profiles across E_R_ binding ratios, we employed bootstrapping sampling of different antibody binding bins (E_R_ of 0.1,1-10,10-100, >100) and sampled across different immunization and antibody profiles. High bootstrap scores indicate robust repertoires sampled at sufficient depths (Supporting Figure 10). Individual mutation logo plots showcased for a representative droplet of each sequenced repertoire further highlight the different profiles across the RBD in full detail (Supporting Figure 11). Cosine similarity heatmaps were generated across a matrix of droplets within each repertoire.

### Structural data analysis

To structurally visualize the enriched positions in the three-dimensional context, the cumulative enrichment ratios for each non-synonymous amino acid position was used. Enrichment ratios were normalized before being converted into a RGB value ranging from grey [0.5,0.5,0.5], corresponding to a low enrichment ratio, to red [1,0,0], corresponding to a high enrichment ratio. The normalized colors are projected onto a crystal structure of the RBD in contact with ACE2 (PDB: 6M0J)^39^. This presents a structural overview of the enriched positions and can be used to help identify antibody epitopes and variants that are mutated in the interface between RBD and ACE2.

### Principle Component Analysis (PCA)

PCA was performed using scikit-learn^40^ to reduce multidimensional complexity of the antibody repertoire data. Input for the PCA was assembled from single-cell/single-barcode data, defined as E_R_ value features for each barcode-immunization/condition pair. Each dot in the PCA plot represents a droplet barcode (single antibody). Points were annotated by immunization type and overlap was limited by introducing minimal gaussian jitter (σ= 0.1). PCA analysis was performed in python (Seaborn) using a separate Jupyter Notebook^41,42^.

### Statistical analysis

If not stated otherwise, replicates are presented as mean ± SD, and n refers to the number of biological replicates. Differences in distributions were assessed using two-sided, unpaired, non-parametric Kolmogorov-Smirnov (KS) tests with 95% confidence. To compare epitopes in the RBDs, regions of 10 AA were chosen (Figure 2D), and pairwise comparison between immunizations was performed using one-way ANOVA on ranks by Kruskal-Wallis test (KS). For qPCR comparison (Figure 1C), ordinary one-way ANOVA was used. For comparison of distribution of secretion rate and antigen relocation, KS tests were performed. p-values were denoted as follows: * (0.05 – 0.01), ** (0.01 – 0.001), *** (0.001 – 0.0001), **** (< 0.0001).

## Notes

https://github.com/lucaSCHLTBER/SingleAntibodyDMS/

### Competing Interest Statement

The authors have declared no competing interest.

