## Supplementary Data and Figures for "Single-antibody mutational scanning reveals poly-specificity and immune constraints in vaccination responses"

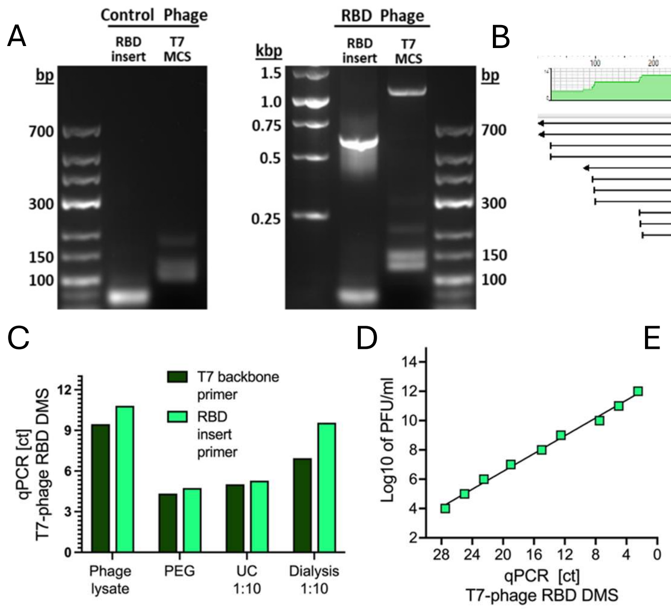


| Purification stage | RBD Phage library (PFU/ml) | |
|---|---|---|
| | Stock 1 | Stock 2 |
| Phage Lysate | $1.6x10^{10}$ | $3.2x10^{10}$ |
| PEG | $3.1x10^{11}$ | $5x10^{11}$ |
| UC | $1.2x10^{12}$ | $0.8x10^{12}$ |
| Dialysis | $9.9x10^{10}$ | $9.2x10^{10}$ |

**Supporting Figure 1.** Cloning, generation and characterization of T7-RBD_DMS antigen library. **A)** PCR of the inserted RBD DMS library. The receptor binding domain deep-mutational scanning library is inserted into a T7 vector backbone using EcoRI and HindIII overhang primers. Insert is verified using primers inside the RBD library (RBD insert) and outside of the T7 Multiple Cloning site (T7 MCS). **B)** Sanger sequencing of the RBD library. Alignment of the T7 insert from the amplified library against the SARS-CoV-2 RBD reference sequence (Plasmid Addgene 1000000172). Downstream of the RBD lies gene10 capsid protein of the T7 including the T7-tag. **C)** RBD phage library quantification at different purification stages using T7-MCS specific primer and RBD insert specific primer. **D)** Correlation between PFU/ml (Plaque Assay) and qPCR (ct) for T7 phage titer calculation. **E)** Table with the PFU/ml transformed titer of two replicate T7-RBD phage library stocks (from C) at the different purification steps.

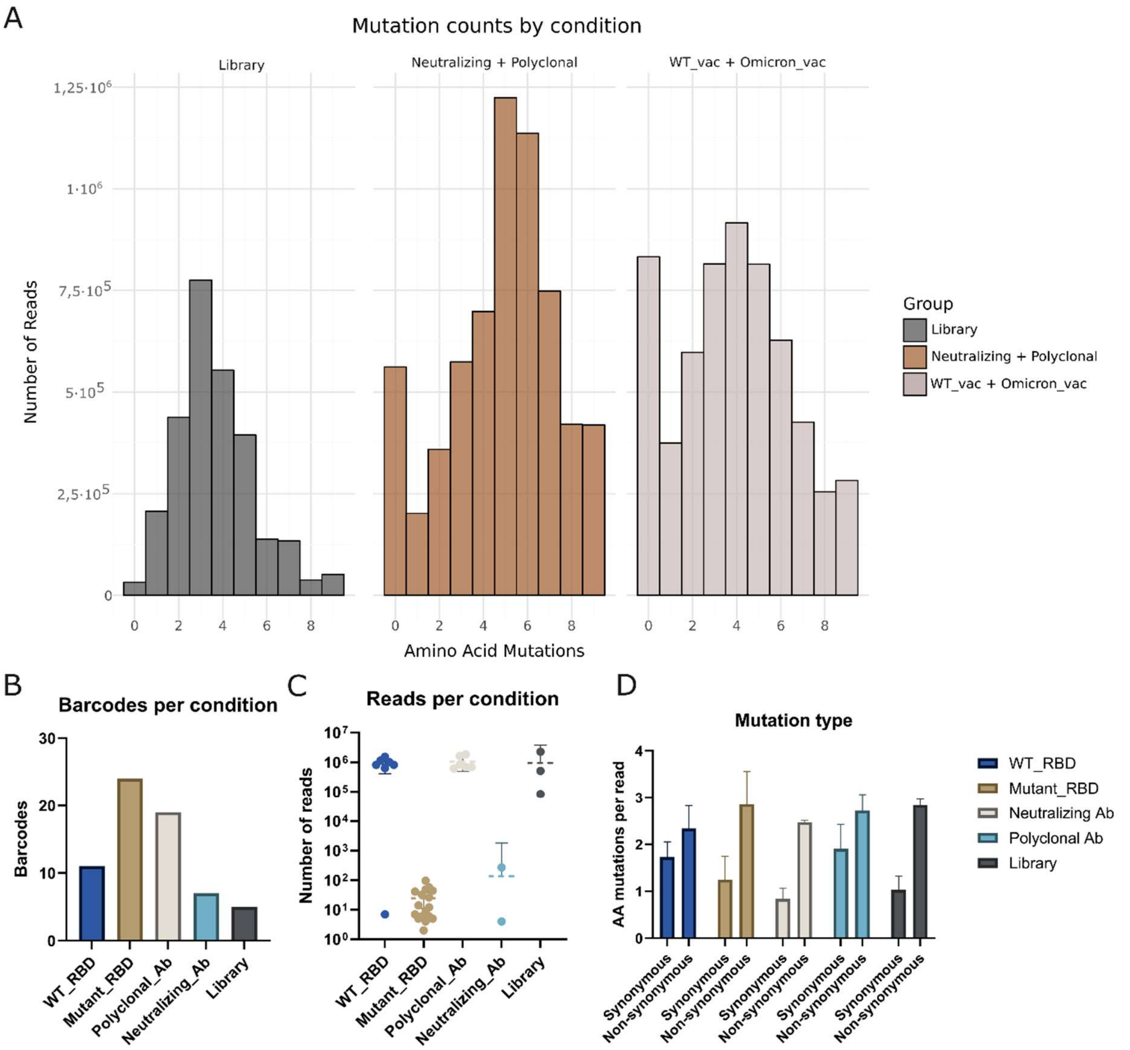


**Supporting Figure 2.** Sequencing characteristics. **A)** Overview of the number of amino-acid mutations per read across different groups. Mutation is based on codon differences. **B**) The number of barcodes retrieved per conditions. **C)** The amount of reads per condition. WT_RBD showed high read counts as compared to the mutant_RBD which generally showed low read counts. **D)** The average amount of synonymous or non-synonymous amino-acid mutations per read. Overall, a higher rate of non-synonymous mutations is seen across all conditions.

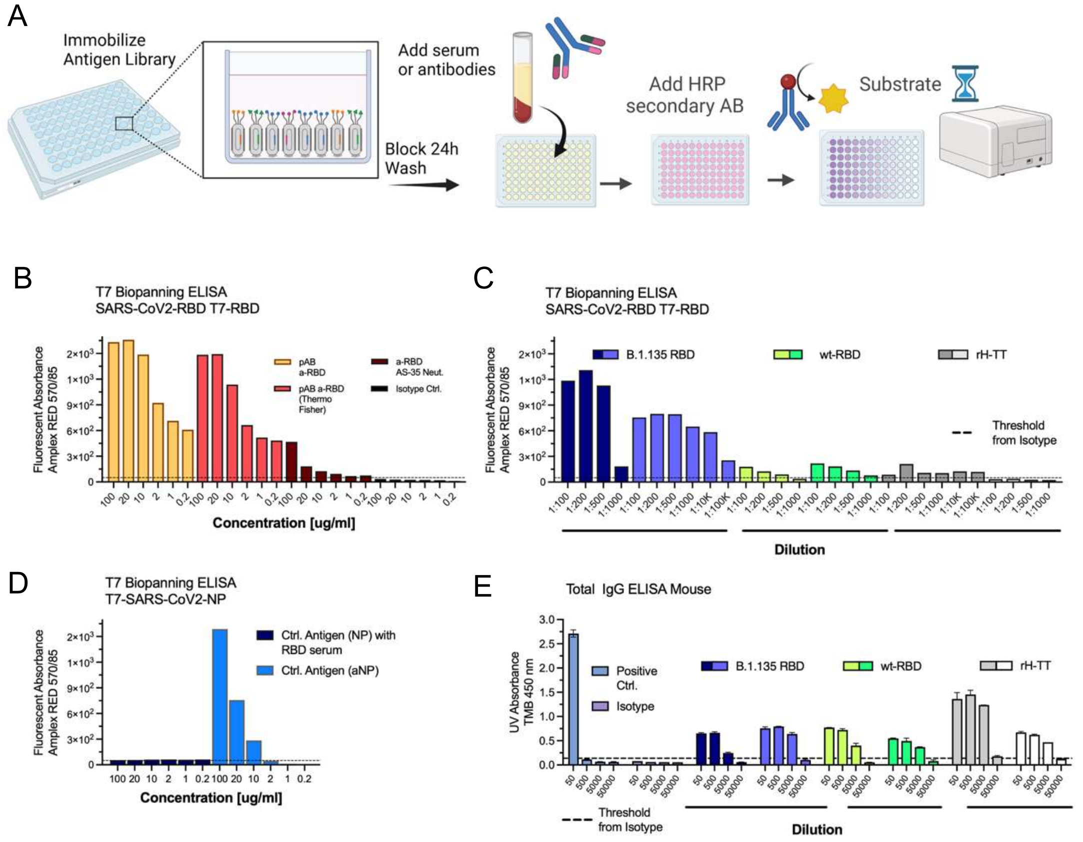


**Supporting Figure 3.** Biopanning ELISA using the purified T7-Receptor-binding-domain deep mutational scanning library confirms binding of calibration antibodies and polyclonal serum. **A)** T7 ELISA Workflow and data from analysis of plasma serum from vaccinated mice, mono-or polyclonal antibodies. For binding assay, first Antigen library is immobilized on bio-panning 96-well plates over-night before incubation with respective sample. **B)** T7-DMS Biopanning cross-validation of calibration antibodies used in single-cell relocation and antigen-scanning experiments. T7-variant binding was measured at different concentration of antibodies measured at 1:2 dilutions starting at 100ug/ml. Antibodies were diluted from original stocks 1 µg/ml. **C)** T7-RBD variant binding polyclonal sera were serial diluted from 1:100 starting dilution. Plotted are individual measurement from serial dilutions measured in the same plate. **D)** T7 bacteriophage expressing SARS-CoV-2 NP does not bind RBD immunized serum, but binds anti-NP antibody. T7-variant binding was performed at different concentration of antibodies measured at 1:2 dilutions starting at 100ug/ml. Antibodies were diluted from original stocks 1 µg/ml. **E)** Antibody level determination in immunized mouse sera via direct ELISA. Positive control, isotype as well as individual can be determined via color legend, immunization groups via horizontal label. Limit of detection is indicated as dashed black line. Positive IgG signals could be detected for all mouse sera and were measurable up to dilutions of 1:5,000 and 1:50,000. No significant differences in antibody levels were found between the different immunization groups. For 3C and 3E, mice were immunized with rH-TT: recombinant-heavy chain tetanus toxin; wt-RBD: wild-type (Wuhan) Receptor Binding Domain (recombinant protein) or B.1.135 RBD: the Receptor Binding Domain of the SARS-CoV-2 B.1.135 lineage (recombinant protein).

**Supporting Information SI Table 1 – used reagents:**

| Antibodies used in this study: | | Supplier | CAT Number |
|---|---|---|---|
| Capture reagent mouse | CaptureSelect™ Biotin Anti-LC-kappa (Murine) Conjugate | ThermoFisher | 7103152500 |
| Capture reagent human | CaptureSelect™ Biotin Anti-IgG-Fc (Human) Conjugate | ThermoFisher | 7103262100 |
| Anti-Tail T7 | T7 Tailfibre mAb | Merck Millipore | 71530 |
| aSpike | Anti-SARS-CoV-2 Spike antibody | SinoBio | 40591-MM41 |
| nAb | Anti-RBD neutralizing antibody | AcroBiosystems | SAD-S35 |
| IgG detection probe | Goat Anti-Mouse IgG Fc AF647 (Goat) | Southern Biotech | 1033-31 |
| IgG detection probe | Goat Anti-Human IgG Fc AF647 (Goat) | Southern Biotech | 2048-31 |
| IgG detection probe | Goat Anti-Mouse IgG Fc AF488 (Goat) | Southern Biotech | 1033-30 |
| Isotype Control DT antibody | Diphtheria Toxin monoclonal antibody G97E | ThermoFisher | MA1-7028 |
| T7-PE | Rabbit Anti T7-Tag antibody, PE-labelled | Abcam | ab72563 |
| C1 | Anti-RBD antibody C1, clone H4 | Invivogen | cov2rbdc1-mab10-3 |
| C2 | Anti-RBD antibody C2, clone B38 | Invivogen | cov2rbdc2-mab10-3 |
| E1 | Anti-RBD antibody e1, clone CR3022 | Invivogen | srbd-mab10-3 |
| pAb | Anti (RBD) polyclonal antibody (mouse) | Proteogenix | PTXCOV-A539 |
| pAb II | Anti (RBD) polyclonal antibody (rabbit) | Proteogenix | PTXCOV-A536 |
| aCD20 | Anti-CD20 Antibody L26 (ELISA isotype Ctrl.) Mouse Monoclonal IgG2a | Antibodies.com | A253774 |
| wt-RBD (RBD Wuhan) | RBD for immunization 1 - SARS-CoV-2 S protein RBD, His Tag(SPD-C52H3) is expressed from human 293 cells (HEK293). It contains AA Arg 319 - Lys 537 | AcroBIo | SPD-C52H3 |
| RBD B1.1.135 mutant | RBD for immunization 2 - SARS-CoV-2 S protein RBD (K417N, E484K, N501Y), His Tag (SPD-C52Hp) is expressed from human 293 cells (HEK293). It contains AA Arg 319 - Lys 537 | AcroBIo | SPD-C52Hp |

| Name | 5' Sequence | Supplier |
|---|---|---|
| 1st round PCR Fwd primer EcoRI | TCG TCG GCA GCG TCA GAT GTG TAT AAG AGA CAG GAG GGT CGG CTA GCC ATA TG | Microsynth |
| 1st round PCR Rev primer HindIII | GTC TCG TGG GCT CGG AGA TGT GTA TAA GAG ACA GTT CGG AAC CGC CCC CCT CGA G | Microsynth |
| T7 MCS primer (T7 UP & T7 DOWN) | GGAGCTGTCGTATTCCAGTC/ AACCCCTCAAGACCCGTTTA | Novagen T7 Select |
| qPCR RBD (140bp) | CGGTTCTCACCCCTCAACAA/ AGCCTCCTCCACCACTAGCC | IDT |
| qPCR T7 (141bp) | ATGGTGCGGTTCTGGCTGAG/ TACCCAGCGCAACTTGGTCG | IDT |
| AmpliconSeq Primers for DMS-RBD library sequencing FWD | TCG TCG GCA GCG TCA GAT GTG TAT AAG AGA CAG GAG GGT CGG CTA GCC ATA TG | Microsynth |
| AmpliconSeq Primers for DMS-RBD library sequencing REV | GTC TCG TGG GCT CGG AGA TGT GTA TAA GAG ACA GTT CGG AAC CGC CCC CCT CGA G | Microsynth |

**Supporting Information SI Table 2 – measured SPR affinities for selected antibodies.** Measured against the full spike protein determined on an SPR-24 Pro (Bruker Daltonics) using a titration cyclie kinetics assay.

| Antibody Clone | *Measured dissocation constant $K_D$* [nM] |
|---|---|
| Anti-RBD antibody C1, clone H4 | 0.55 |
| Anti-RBD antibody C2, clone B38 | 0.92 |
| Anti-RBD antibody e1, clone CR3022 | 0.16 |
| Anti (RBD) polyclonal antibody pAB II* | ~36 |

* Accurate determination of $K_D$ was not possible; most likely due to polyclonal nature, and rabbit polyclonal was used due to discontinuation of the murine product.

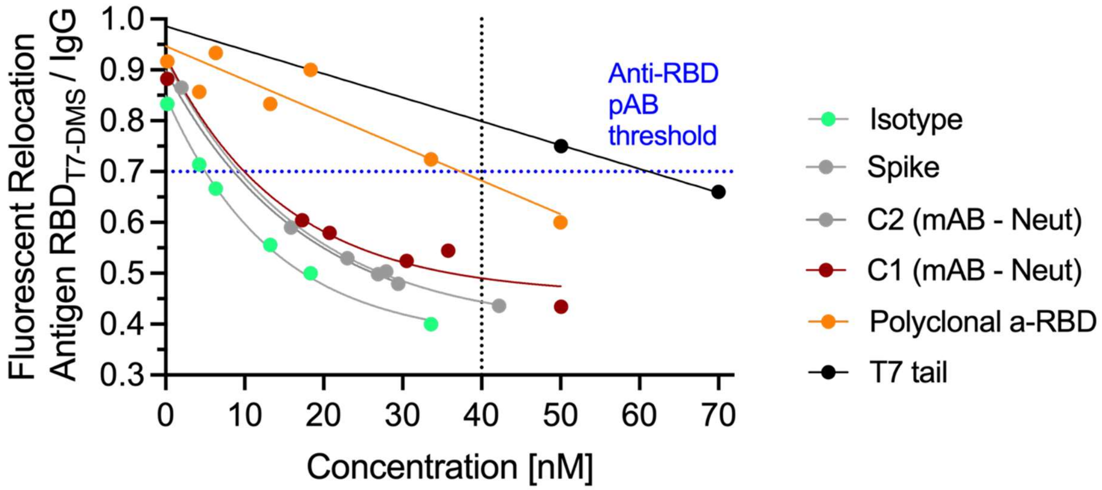


**Supporting Figure 4.** Calibration of the developed T7 assay for IgG secretion in stationary droplet arrays of encapsulated antibody as a surrogate for anti-RBD cross-reactivity. Droplets are encapsulated with murine IgG detection assay, nanoparticles, T7-RBD library and anti-T7-tag PE probe at different concentrations [nM]. All Isotypes are IgG. Beadline relocation (R) is calculated from fluorescent intensities onto nanoparticles relative to droplet background of the respective channel and are calculated using the DropMap script implemented from Matlab [1]. Dotted line indicates in-droplet antibody concentration used for subsequent stratification of mono- and poly-reactive antibody-secreting cells.

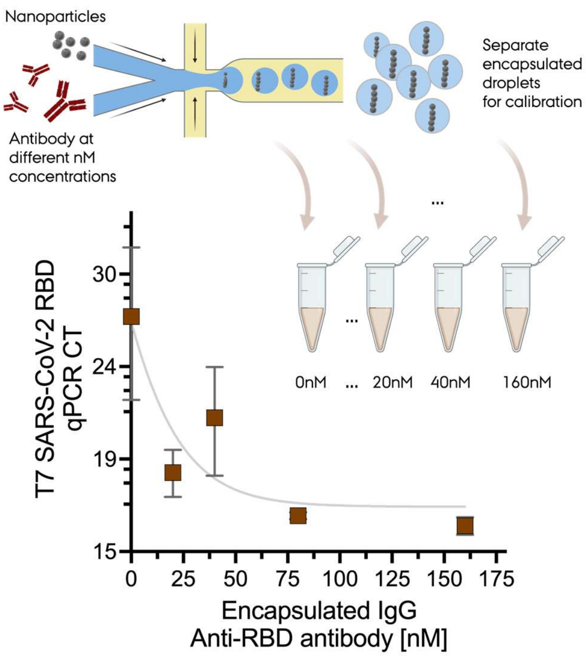


**Suppoirting Figure 5.** Correlation between antigen-qPCR $C_t$ values and the concentration of added antibody, in this case anti-SARS-CoV-2 RBD pAb. The data was acquired from 1,000 dememulsified droplets for the number of recognized RBD-expressing T7 phages. The plot shows the average Log10 $C_t$ and standard deviation. The data displayed a decrease in $C_t$ when the concentration of antibody was increased as expected (Fit was computed using a one-phase association fit in Prism).

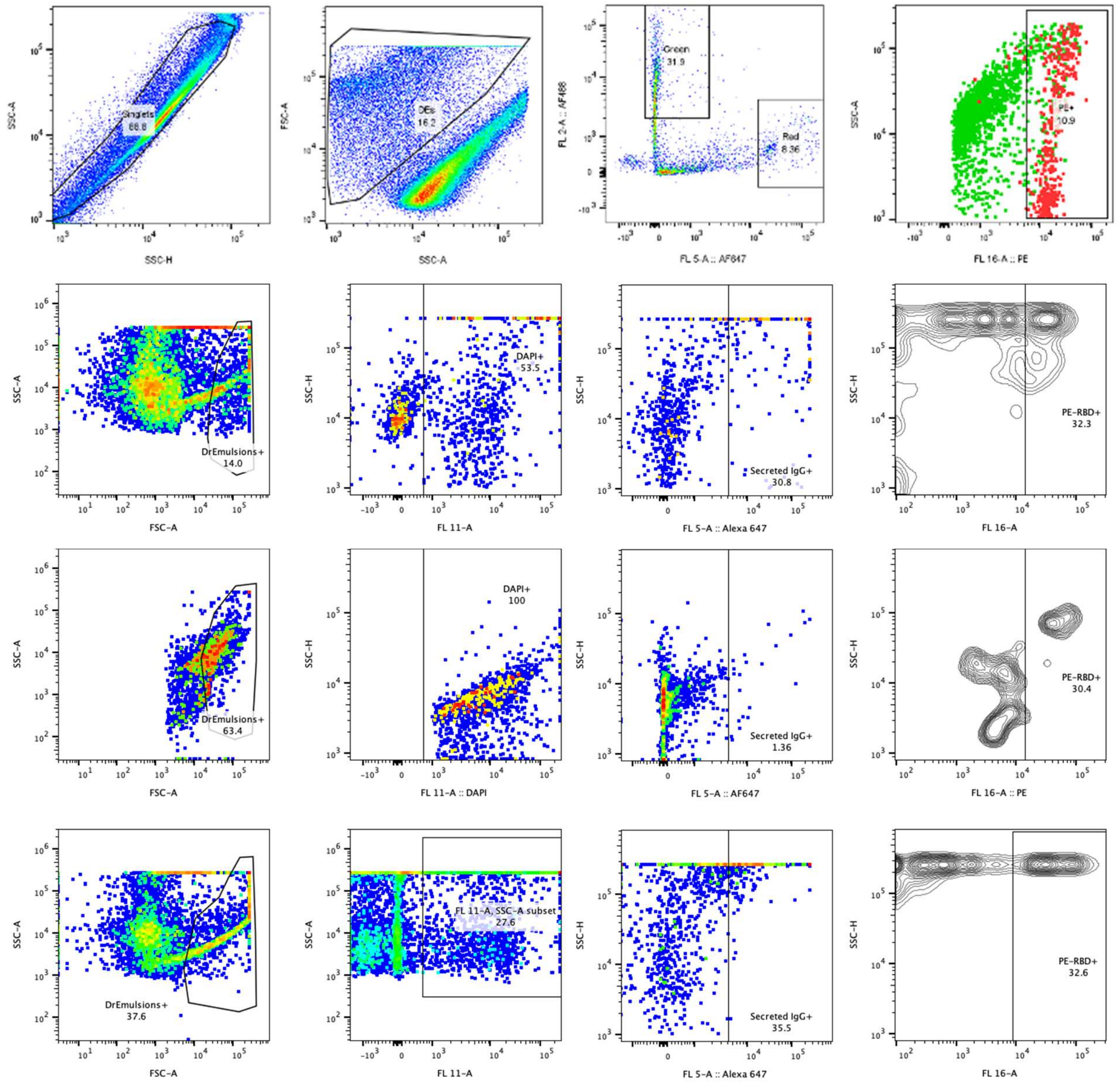


**Supporting Figure 6.** Gatting strategy for B cells from immunization studies for single droplet sorting (FADS). Droplets were encapsulated with isolated, DAPI stained (CTV – Cell proliferation kit), murine B cells (inlet 1) and the T7-RBD-DMS library, nanoparticles and an anti-IgG-FC AF647 (Southern Biotech) detection probe (inlet 2). Double Emulsion droplets were generated as described in Methods and were gated by FSC/SSC (gating was optimized on each produced emulsion), selected by DAPI and IgG-secretion (DAPI$^{+}$/AF647$^{+}$). Finally, populations were gated and sorted based on their RBD-Variant PE signal from an anti-T7tag detection probe (Goat). For calibration, the anti-SARS-CoV-2 RBD polyclonal AB (Proteogenix) was used (top); for antigen-scanning from individual B cells, three example sample for single droplet sorting are displayed (Bottom). Plots were generated with FlowJo 10.10.0.

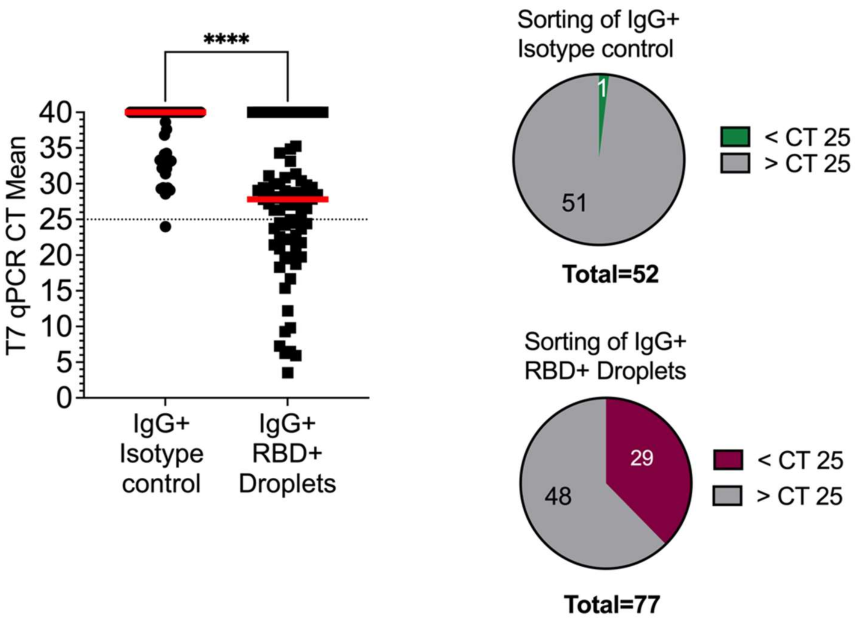


**Supporting Figure 7**. qPCR evaluation of Droplet Sorting Efficiency (post-sorting). Individual droplets from positive events (anti-RBD pAB) and negative events (Isotype control) were sorted into 96-well PCR plates using ThermoFIsher BigFoot Sorter. Antigen binders were washed on a magnetic rack three times before lysing T7 and subsequent qPCR for T7 vector arms.

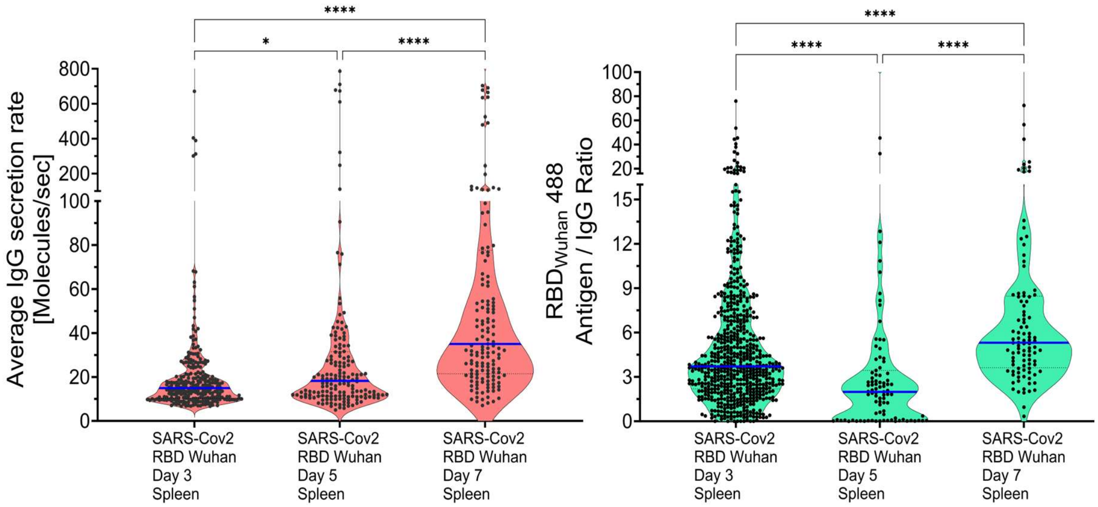


**Supporting Figure 8.** Secretion rate and antigen binding (RBD monomer) of individual IgG-secreting B cells after immunization with SARS-CoV-2 RBD Wuhan at different days post 2$^{nd}$ immunization. Data from stationary droplet array for IgG secretion. **Left:** Average secretion rate (in IgG molecules per second) of individual isolated splenic B cells from RBD-WT immunized mice on day 3, 5 and 7 post-secondary immunization. Displayed is the secretion rate of IgG calculated from the relocalization of an anti-IgG-647 detection probe upon secretion of IgG on to VHH-IgG nanoparticles for each droplet containing an individual murine B cells. The secretion rate was calculated and converted from a calibration curve using a murine IgG isotype control. **Right:** Antigen density as the ratio of relocation of antigen relative to the secreted IgG over time. Antigen density was calculated from the relocalization of an AF488 fluorescent-labeled recombinant RBD (Wuhan) protein on to IgG capture nanoparticles at each time-point of measurement. Recombinant, labeled RBD was incorporated into the droplet assay (instead of T7-RBD-DMS). Statistics were performed using ONE-Way Anova on ranks Kruskal-Wallis test (KS). p-values were denoted as follows: * (0.05 – 0.01), ** (0.01 – 0.001), *** (0.001 – 0.0001), **** (< 0.0001). Medians are displayed as blue lines.

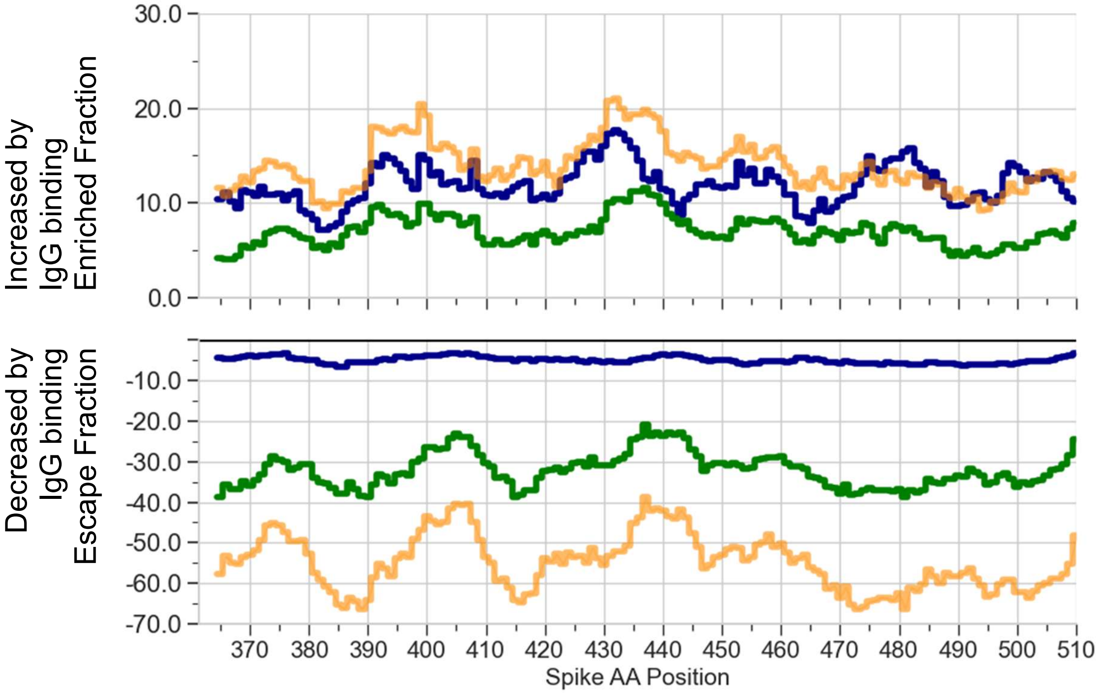


**Supporting Figure 9.** Sum of enrichment ratios $E_R$ (Log10 change of mutation frequency) of B.1.351 immunization (blue), wt-RBD (green) and pAb (orange) from all IgG+ poly-specific antibodies split by enriched (top) and escape (bottom) fraction. X-axis represents Spike Amino acid position in the receptor binding domain (RBD).

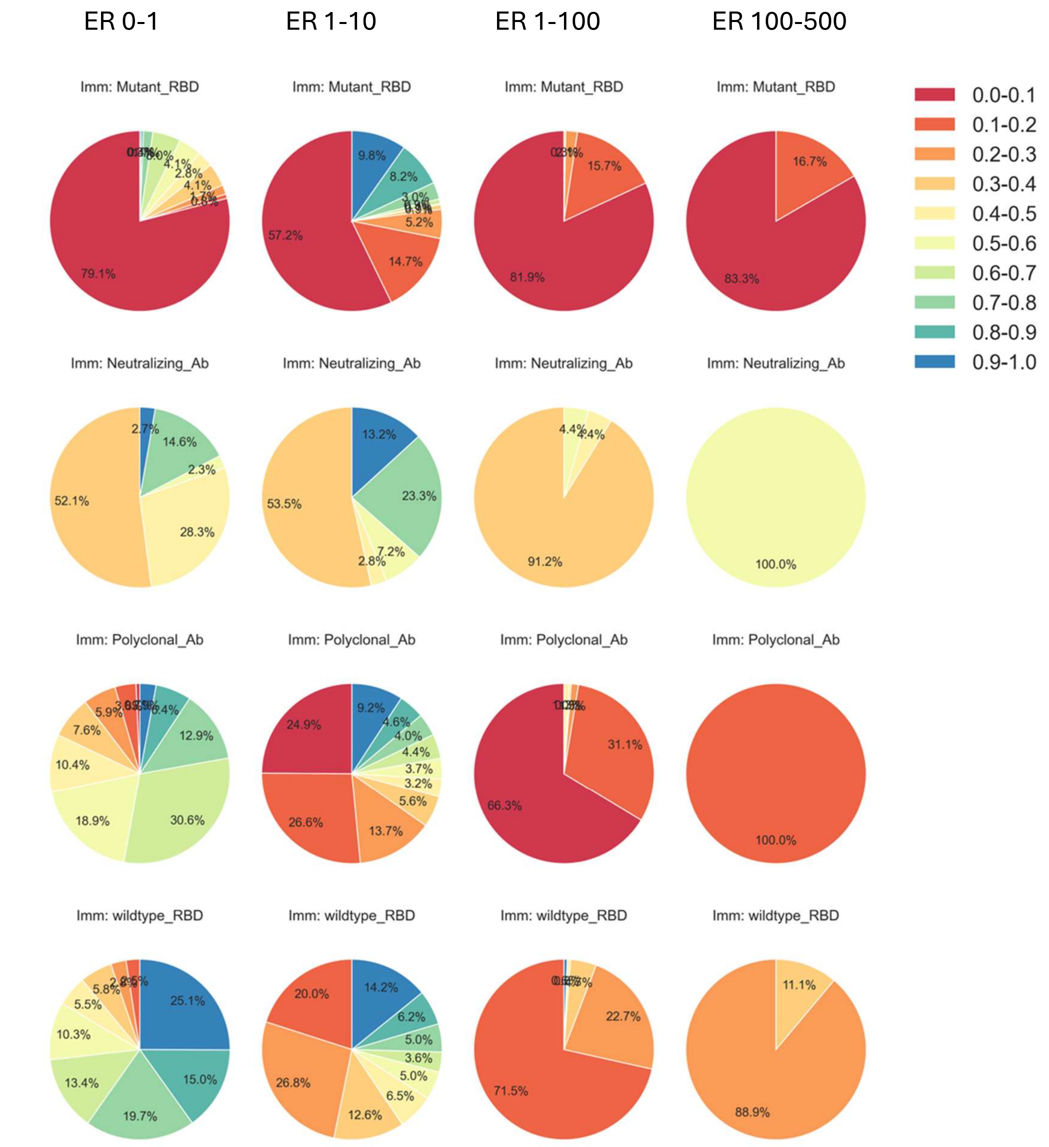


**Supporting Figure 10.** Pie charts illustrating the distribution of bootstrap mean frequency (Antibody binding) categorized by immunization type and enrichment bin. Each row corresponds to a distinct immunization group, while each column represents a specific enrichment bin. Pie charts are separated according to Enrichment Ratio such as 0-1 (immune escape), 1-10 ( up to 10x fold increase in binding), 10-100 (up to 100-fold increase in binding), 100-500 (up to 500-fold increase in binding). Within each pie chart, the proportion of samples falling into bootstrap mean frequency bins ranging from 0.0 to 1.0 (in increments of 0.1) are displayed as a metric for the degree of mutation variant binding across the single-droplet repertoire (1.0 = homogeneously distributed, 0.1 mutation which is very rarely relevant in the immune repertoire). Different frequency ranges are color-coded using a consistent spectral palette, with percentages labeled on each pie segment to indicate their relative contribution. Empty plots indicate combinations of immunization and enrichment bins with no data. The title above each pie chart specifies the immunization and enrichment bin it represents.

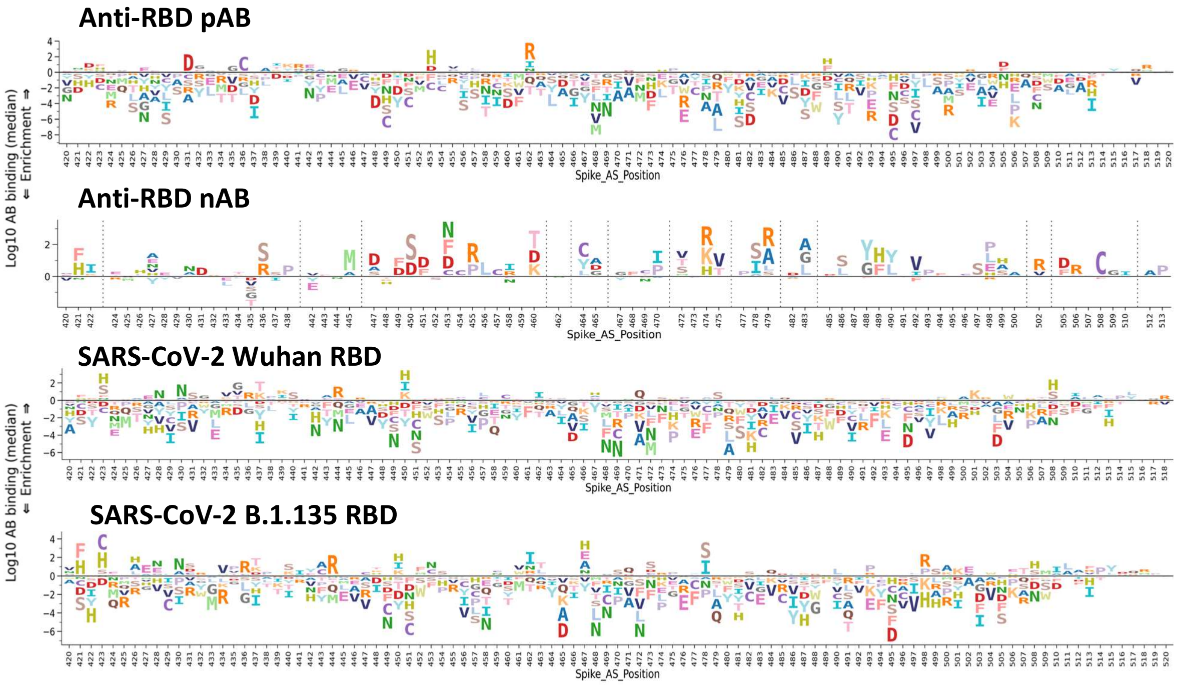


**Supporting Figure 11.** Representative, example (single-droplet) logo plots for mutational variants increased in frequency by antibody binding (Binding Fraction, >0) and decreased (Escape fraction <0). Y-axis plots the median (log10) IgG binding ratio of the SARS-CoV-2 RBD variants across Spike protein position 420-520. Logoplots are colored by amino-acid. Graph generated with DMSlogo package [2].

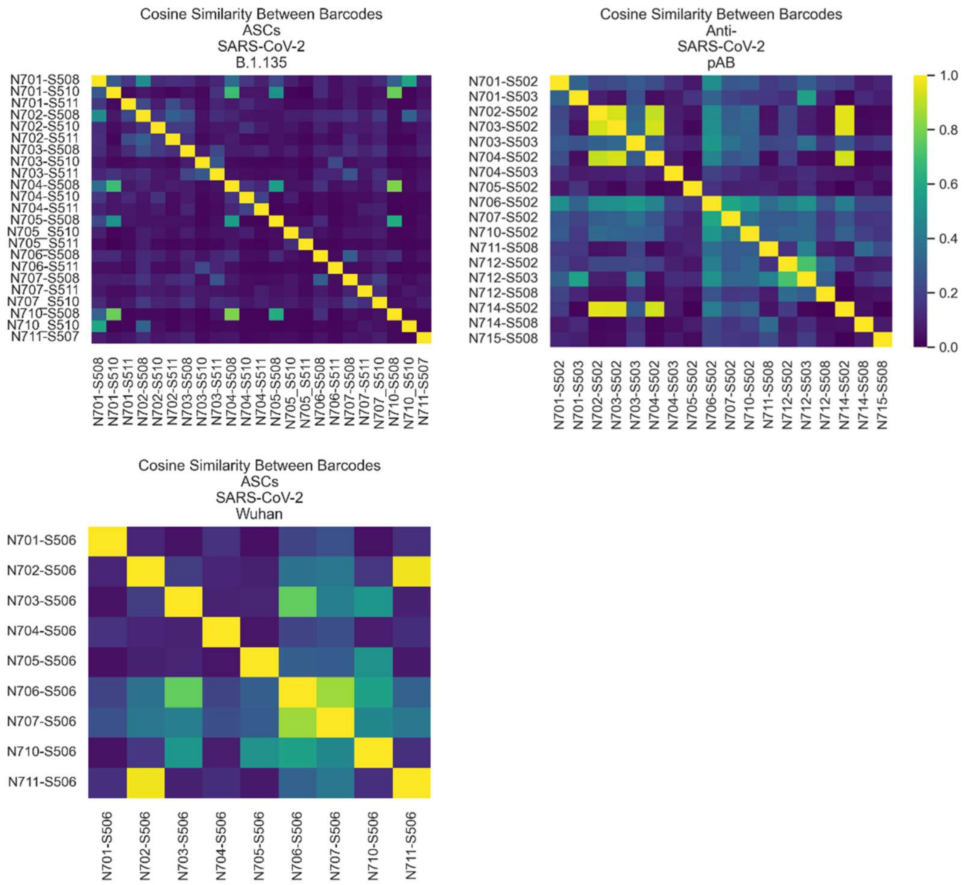


**Supporting Figure 12. Pairwise cosine similarity heatmap between barcodes of immunization group.** Each cell in the matrix represents the cosine similarity between two barcodes' enrichment profiles across all tested amino acid substitutions at spike positions. Higher similarity values (closer to 1, yellow) indicate more consistent epitope recognition patterns, whereas lower values (blue) reflect greater divergence among antibody specificities within this immunization group. Cosine similarity between individual droplet barcodes (Antibody-secreting cell Variant profile) of B.1.135 RBD (**top-right**) and (pAB) polyclonal anti-SARS-CoV-2 antibody (**top-right**) and wt-RBD (Wuhan) immunized mice (**bottom**). The number of sequenced cells and antibodies for the analysis include B.1.135 RBD (n=23), pAB (n=18) and wt-RBD (n=9), totaling 50 sequenced events.

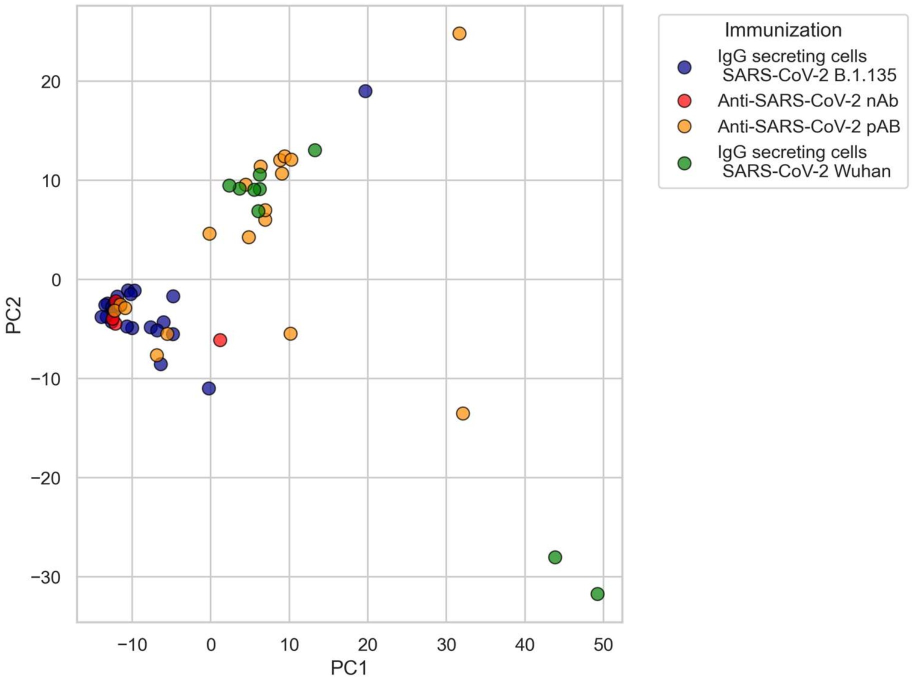


**Supporting Figure 13.** Visualization of SARS-Cov-2 variant antibody enrichment profiles from immunizations and calibration antibody data using PCA global dimensionality reduction. Each dimension is defined as the enrichment ratio for a specific Amino acid mutation within the SARS-CoV-2 Spike protein sequence (RBD only). Each individual point represents a unique droplet barcode and is colored by its immunization condition.

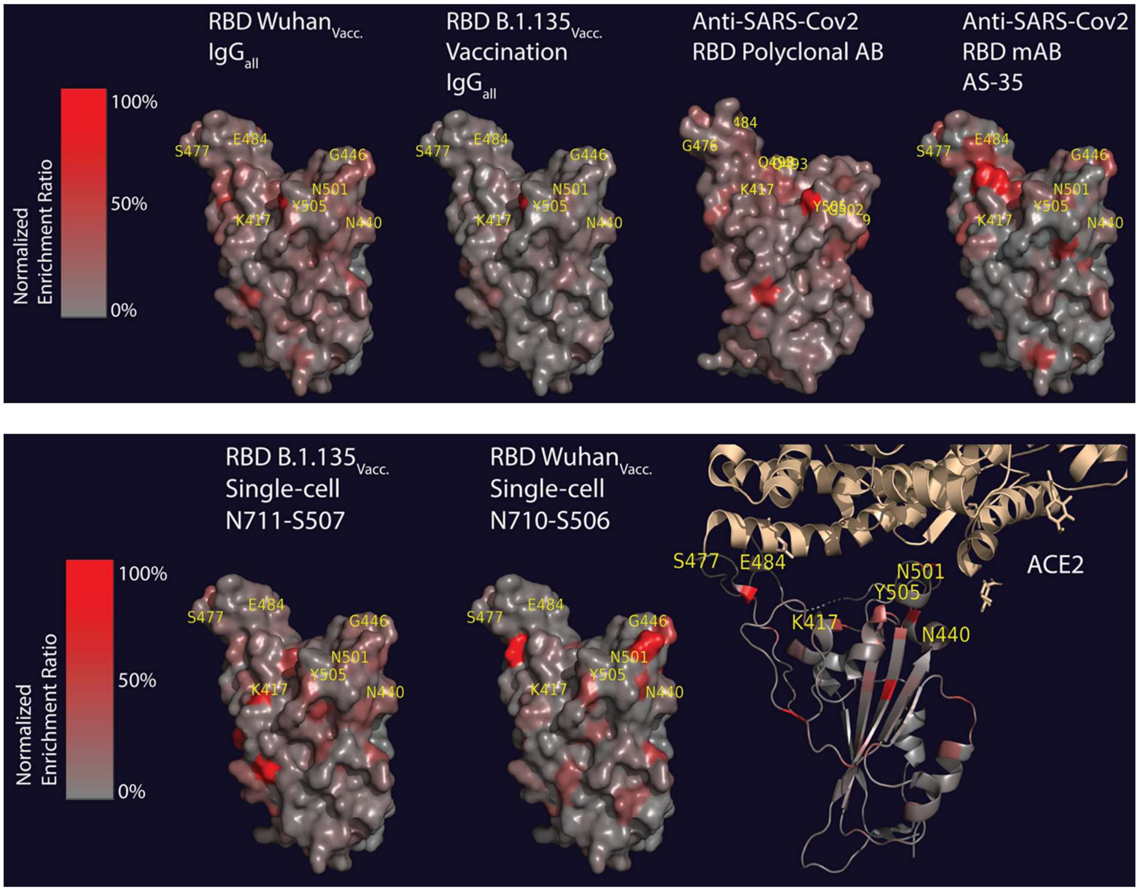


**Supporting Figure 14:** Top: Structural representation of contact sites of grouped antibody enriched variants from I. secreted mAB from RBD Wuhan vaccinated mice, II. secreted mAB from RBD B.1.135 vaccinated mice, III. anti-SARS-CoV-2 RBD polyclonal AB and IV. monoclonal, neutralizing antibody clone AS-35. SARS- CoV-2 Wuhan SPIKE RBD. Bottom: Single-cell antibody variant profiles from IgG secreting cells from RBD Wuhan and RBD B.1.135 vaccination derived antibodies. SARS-CoV-2 Wuhan SPIKE RBD structure were imported using Pymol and and colored based on the enrichment matrix for each spike position normalized to 100%. PDB= 6m0j

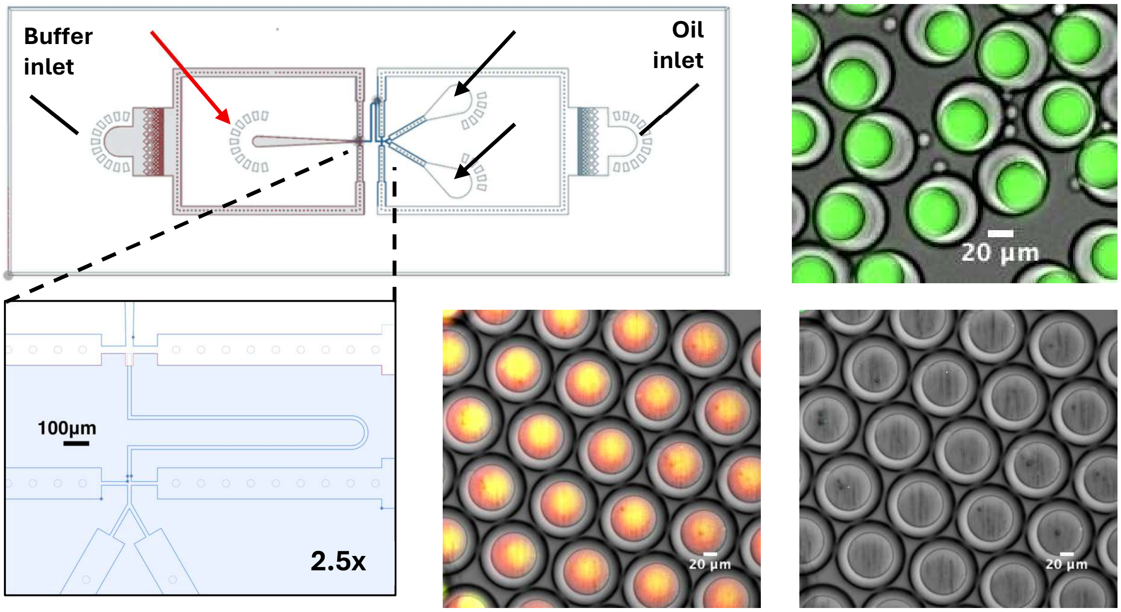


**Supporting Figure 15:** Design of the adapted double emulsion pico-reactor microfluidic chip for Fluorescent Activated Droplet Sorting (FADS): Oil inlet provides the outer layer for the 1$^{st}$ (W/O) droplet encapsulation of nanoparticles and cells (inlet 1 and 2, indicated by black arrows). Droplets are led through a transition channel (shown enlarged below) and subsequently W/O droplets are encapsulated again into (hydrophilic W/O/W) aqueous droplets. Outlet for collecting droplets for sorting is indicated by a red arrow. Original design adapted from Brower *et al*. [3]. Right side shows example double-emulsion droplets encapsulated with Calcein-Green (**top**) and double-emulsions droplets contain Antibody Secreting Cells, T7 Antigen Libraries and nanoparticles after encapsulation (**bottom**).
